## Supplementary Figures for "Ultra-mild bisulfite outperforms existing methods for 5-methylcytosine detection with low input DNA"

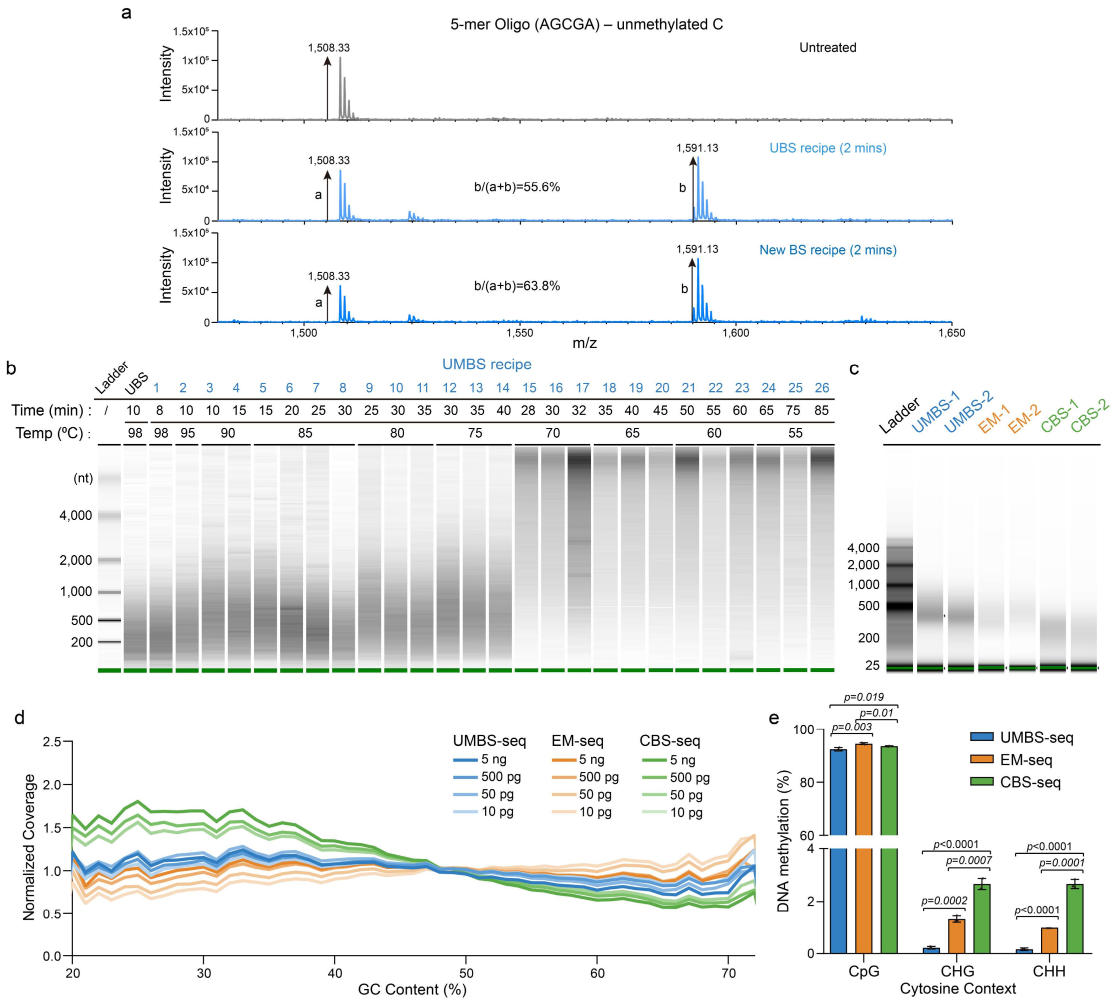


**Supplementary Fig. 1: Development of the Ultra-Mild Bisulfite (UMBS) recipe and optimization of UMBS reaction conditions. a,** MALDI-TOF MS analysis of unmethylated 5-mer DNA oligonucleotide (AGCGA) treated with Ultra-fast Bisulfite (UBS) and the newly developed bisulfite recipe for 2 minutes. The relative abundances of the unreacted oligo and the U-BS product indicate that the new bisulfite recipe achieves a higher C-to-U conversion efficiency compared to the UBS recipe. **b,** Comparative analysis of DNA damage under varying UMBS reaction conditions, assessed by bioanalyzer (Agilent RNA 6000 Pico). Intact lambda DNA was treated with the UMBS conversion method across a range of temperatures (55 °C to 98 °C) and reaction times (8 to 85 minutes). **c,** TapeStation electropherograms of fragmented, adapter-ligated lambda DNA processed using the three indicated methyl conversion methods. **d,** Normalized GC coverage distribution across the lambda phage genome. Libraries prepared using UMBS-seq and EM-seq (NEB) show significantly more uniform GC coverage compared to those prepared with CBS-seq. **e,** Estimated average DNA methylation levels at CpG sites in methylated pUC19 DNA. UMBS-seq was compared with EM-seq and CBS-seq as benchmarks. Statistical significance was evaluated using two-tailed Student's t test; p-values are indicated. N = 3 libraries per method prepared from the 5ng DNA input were analyzed. Data are presented as mean values +/- SD.

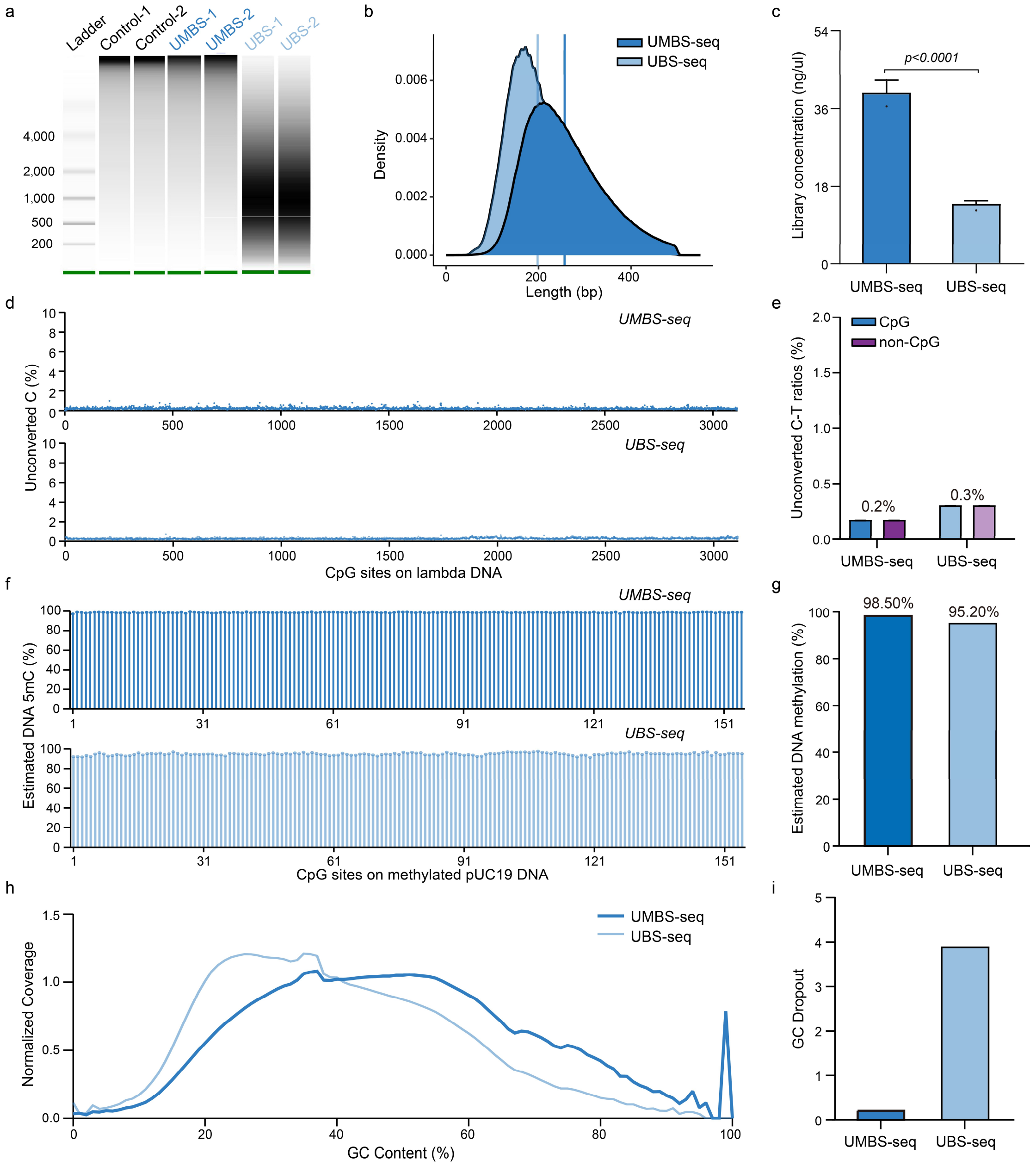


**Supplementary Fig. 2: Systematic performance comparison between Ultra-Mild Bisulfite (UMBS) and Ultra-fast Bisulfite (UBS) methyl conversion methods. a,** Comparative analysis of DNA damage in 100 ng of intact lambda DNA induced by untreated, UMBS, UBS methyl conversion methods, assessed using a bioanalyzer (Agilent RNA 6000 Pico). **b,** DNA insert size distribution of sequencing libraries prepared from 10 ng mESC gDNA, demonstrating that UMBS-seq preserves longer DNA insert size with reduced DNA damage than UBS-seq. **c,** Concentrations of sequencing libraries generated from the specified lambda DNA amounts after treatment with UMBS and UBS conversion methods. Library concentration was measured after 9 PCR cycles with a 20 µL final elution volume. Statistical significance was assessed using two-tailed Student's t test; p-values are indicated. N = 2 libraries per method prepared from the 10 ng lambda DNA input were analyzed. Data are presented as mean values +/- SD. **d,** Scatterplot showing the unconverted C ratios at individual CpG site of lambda DNA treated with UMBS and UBS conversion methods. **e,** Bar plot shows the averaged unconverted C-T ratios of cytosines in both CpG and non-CpG contexts in lambda DNA, treated with the UMBS-seq and UBS-seq methods. **f,** Cleveland dot plot showing the estimated DNA methylation levels at each CpG site of methylated pUC19 DNA treated with the UMBS-seq and UBS-seq methods. **g,** Bar plot showing the average DNA methylation levels across all CpG sites in methylated pUC19 DNA treated with the UMBS-seq and UBS-seq methods. **h,** GC-bias plot for UMBS-seq and UBS-seq libraries prepared from 10 ng of mESC gDNA. UMBS-seq shows better GC coverage uniformity compared to UBS-seq. **i,** Bar plot showing the GC dropout ratios in UMBS-seq and UBS-seq libraries prepared using mESC gDNA.

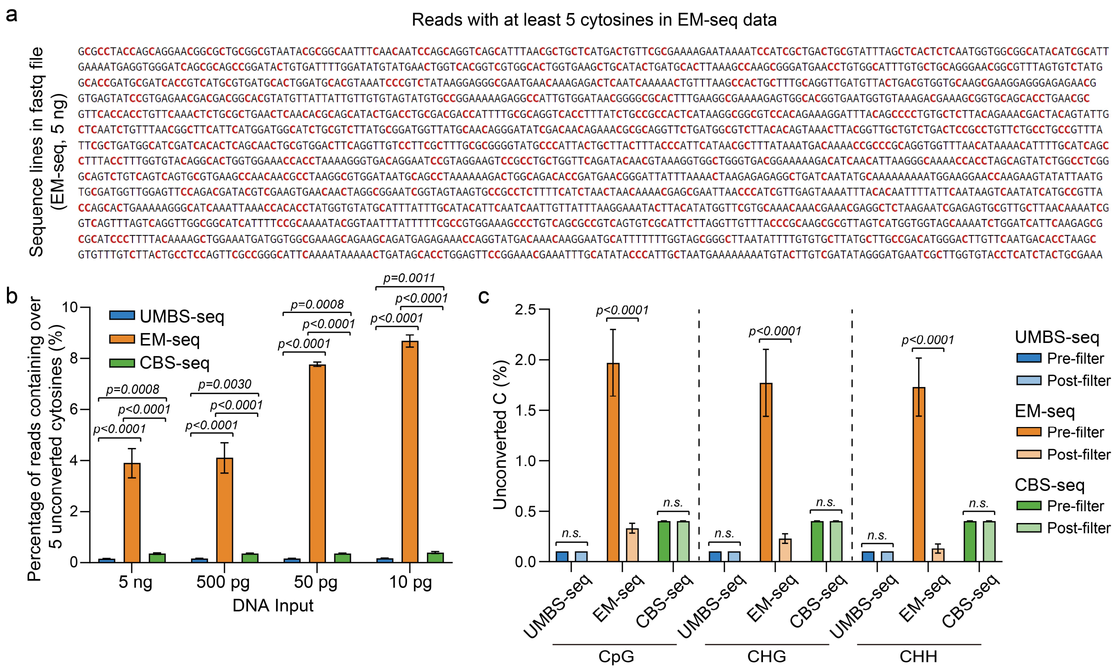


**Supplementary Fig. 3: Presence of unconverted reads in Enzymatic Methylation conversion (EM-seq) libraries. a**, Screenshot of filtered unconverted reads (containing more than 5 unconverted cytosines) from an EM-seq-treated unmethylated lambda DNA library. Unconverted cytosines are highlighted in red. **b**, Bar plot showing the percentage of reads containing more than 5 unconverted cytosines in lambda DNA libraries prepared with each input and method. **c**, Ratios of unconverted cytosines in lambda DNA libraries pre- and post-filtering of reads containing more than 5 unconverted cytosines. Statistical significance was evaluated using two-tailed Student's t test; p-values are indicated. N = 3 libraries per method, prepared from the same DNA input, were analyzed. Data are presented as mean values +/- SD.

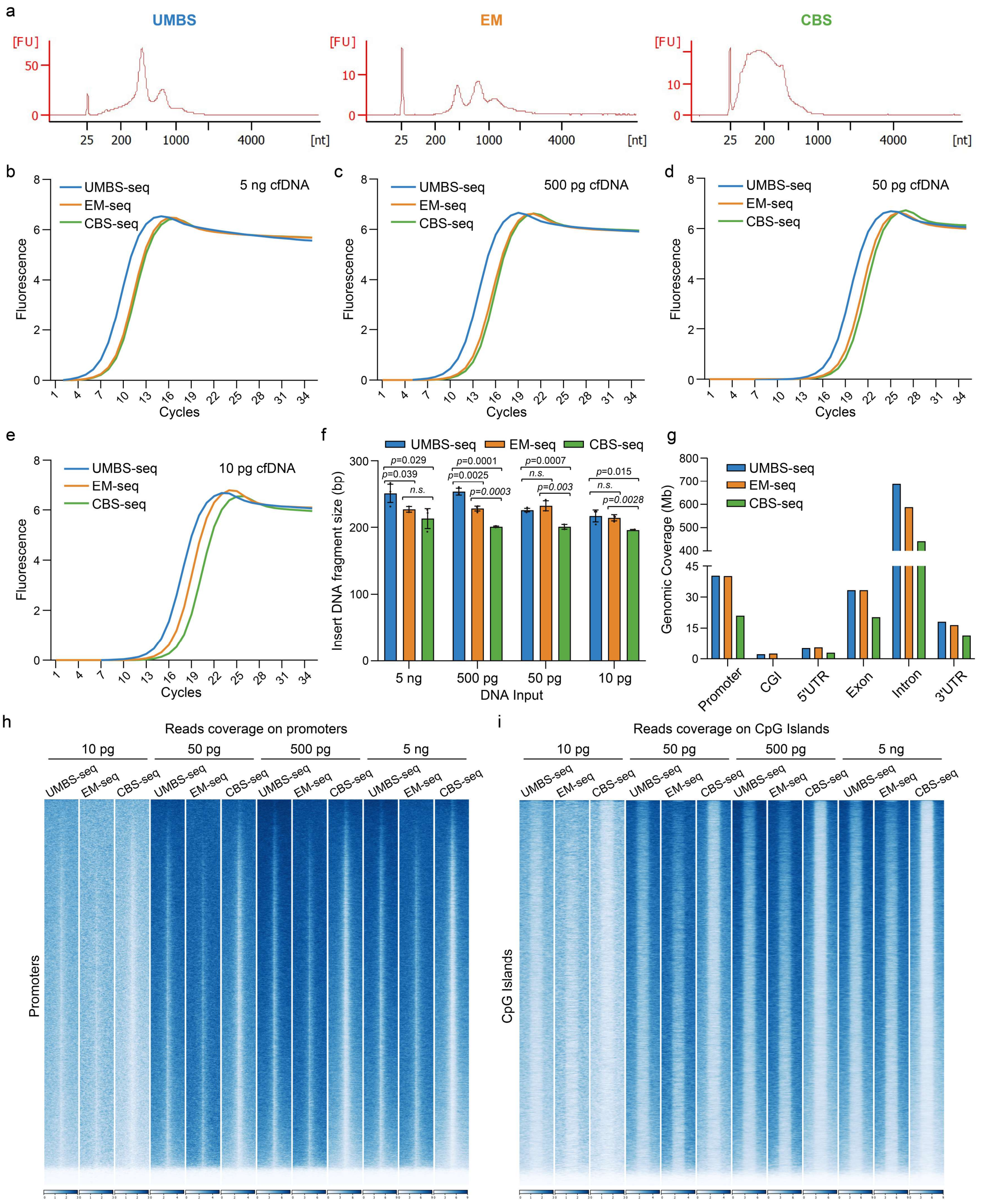


**Supplementary Fig. 4: UMBS-seq demonstrates superior performance on low-input cfDNA. a,** Electropherogram plots of cfDNA following treatment with the UMBS, EM, and CBS methyl conversion methods. **b-e**, Amplification curves from real-time PCR quantification of cfDNA libraries prepared with 5 ng (**b**), 0.5 ng (**c**), 0.05 ng (**d**), and 0.01 ng (**e**) of cfDNA input using UMBS-seq, EM-seq, and CBS-seq methods. **f**, Bar plot showing the averaged DNA insert size of cfDNA libraries prepared using each conversion method at each input. **g**, Genomic coverage (≥1X) of cfDNA libraries across various genomic features. N=3 Libraries were prepared using 5 ng of cfDNA with each method. Data are presented as mean values +/- SD. **h-i**, Heatmaps showing read coverage distribution within a 2 kb window around transcription start sites (TSS) (**h**) and a 1 kb window around CpG islands (**i**). UMBS-seq and EM-seq libraries exhibit greater and more uniform coverage compared to CBS-seq libraries in both regions.

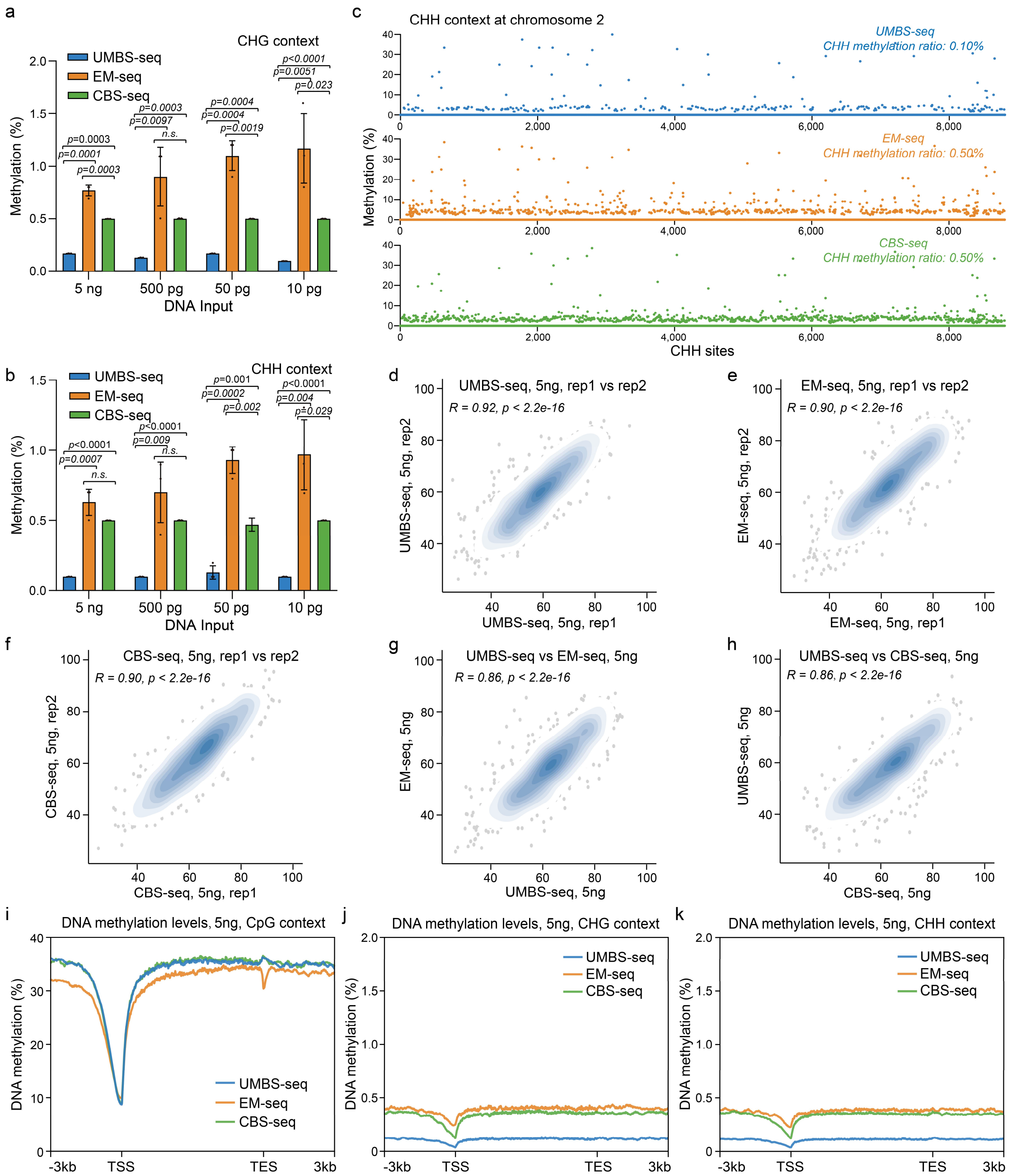


**Supplementary Fig. 5: UMBS-seq exhibits lower background in DNA methylation detection from low-input cfDNA. a-b**, Estimated average DNA methylation levels at CHG (**a**) and CHH (**b**) contexts in cfDNA libraries prepared with varying cfDNA input amounts. UMBS-seq was compared to EM-seq and CBS-seq as benchmarks. Statistical significance was assessed using two-tailed Student's t test; p-values are indicated. N = 3 libraries per method, prepared from the same DNA input, were analyzed. Data are presented as mean values +/- SD. **c**, Ratios of unconverted cytosines at individual CHH sites (over 10X coverage) on chromosome 2 in cfDNA libraries prepared from 5 ng of input using each method. Average methylation levels across these CHH sites are indicated. **d-f**, Scatter plots showing the correlation of DNA methylation levels at individual CpG sites between technical replicates of UMBS-seq (**d**), EM-seq (**e**), and CBS-seq (**f**) libraries, each prepared from 5 ng of cfDNA input. Correlation coefficients and p-values are indicated. **g-h**, Scatter plots showing the correlation of DNA methylation levels at individual CpG sites in 5 ng cfDNA libraries between UMBS-seq and EM-seq (**g**), and between UMBS-seq and CBS-seq (**h**) libraries. Correlation coefficients and p-values are indicated. **i-k**, Metagene profiles comparing average DNA methylation levels across gene bodies in cfDNA libraries prepared from 5 ng of input using each method. Methylation levels were calculated separately for CpG (**i**), CHG (**j**), and CHH (**k**) contexts.

| Performance metrics | UMBS-seq | UBS-seq | EM-seq | CBS-seq |
|---|---|---|---|---|
| Input DNA Range | >=0.01 ng | >=1 ng | >=0.1 ng | >=50 ng |
| DNA Damage | Low | Medium | Low | Hight |
| False Positives | Near Zero | Near Zero | High | Moderate |
| Robustness | Yes | Yes | No | Yes |
| Protocol Time | 2-3 hs | 35 mins | 7-8 hs | 3-4 hs |
| Library Complexity | High | Medium | High | Low |

**Supplementary Fig. 6: Comparison of UMBS-seq with UBS-seq, EM-seq, and CBS-seq based on various performance metrics, including Input DNA Range, DNA damage, False Positives, Robustness, Protocol Time, and Library Complexity.**
